# Leveraging uncertainty information from deep neural networks for disease detection

**DOI:** 10.1101/084210

**Authors:** Christian Leibig, Vaneeda Allken, Murat Seçkin Ayhan, Philipp Berens, Siegfried Wahl

**Author notes:** Co-senior author (Philipp Berens, Siegfried Wahl).

## Abstract

Deep learning (DL) has revolutionized the field of computer vision and image processing. In medical imaging, algorithmic solutions based on DL have been shown to achieve high performance on tasks that previously required medical experts. However, DL-based solutions for disease detection have been proposed without methods to quantify and control their uncertainty in a decision. In contrast, a physician knows whether she is uncertain about a case and will consult more experienced colleagues if needed. Here we evaluate drop-out based Bayesian uncertainty measures for DL in diagnosing diabetic retinopathy (DR) from fundus images and show that it captures uncertainty better than straightforward alternatives. Furthermore, we show that uncertainty informed decision referral can improve diagnostic performance. Experiments across different networks, tasks and datasets show robust generalization. Depending on network capacity and task/dataset difficulty, we surpass 85% sensitivity and 80% specificity as recommended by the NHS when referring 0%–20% of the most uncertain decisions for further inspection. We analyse causes of uncertainty by relating intuitions from 2D visualizations to the high-dimensional image space. While uncertainty is sensitive to clinically relevant cases, sensitivity to unfamiliar data samples is task dependent, but can be rendered more robust.

## Introduction

In recent years, deep neural networks (DNNs)^1^ have revolutionized computer vision^2^ and gained considerable traction in challenging scientific data analysis problems^3^. By stacking layers of linear convolutions with appropriate non-linearities^4^, abstract concepts can be learnt from high-dimensional input alleviating the challenging and time-consuming task of hand-crafting algorithms. Such DNNs are quickly entering the field of medical imaging and diagnosis^5–15^, outperforming state-of-the-art methods at disease detection or allowing one to tackle problems that had previously been out of reach. Applied at scale, such systems could considerably alleviate the workload of physicians by detecting patients at risk from a prescreening examination.

Surprisingly, however, DNN-based solutions for medical applications have so far been suggested without any risk-management. Yet, information about the reliability of automated decisions is a key requirement for them to be integrated into diagnostic systems in the healthcare sector^16^. No matter whether data is short or abundant, difficult diagnostic cases are unavoidable. Therefore, DNNs should report - in addition to the decision - an associated estimate of uncertainty^17^, in particular since some images may be more difficult to analyse and classify than others, both for the clinician and the model, and the transition from “healthy” to “diseased” is not always clear-cut.

Automated systems are typically evaluated by their diagnostic sensitivity, specificity or area under receiver-operating-characteristic (ROC) curve, metrics which measure the overall performance on the test set. However, as a prediction outcome can decide whether a person should be sent for treatment, it is critical to know how confident a model is about each prediction. If we were to know which patients are difficult to diagnose, humans and machines could attend especially to these, potentially increasing the overall performance. In fact, if the machine was making most mistakes when uncertain about a case, one could devise a strategy mimicking typical medical decision making. When faced with a difficult case and feeling uncertain about a decision a junior doctor will consult a more experienced colleague. Likewise, a diagnostic algorithm could flag uncertain cases as requiring particular attention by medical experts.

Estimating the uncertainty about a machine learning based prediction on a single sample requires a distribution over possible outcomes, for which a Bayesian perspective is principled. Bayesian approaches to uncertainty estimation have indeed been proposed to assess the reliability of clinical predictions^19–22^but have not been applied to the large-scale real-world problems that DNNs can target. Outside the medical domain, the integration of the Bayesian ideas and DNNs is an active topic of research^23–34^, but the practical value of the developed methods has yet to be demonstrated.

Due to its ease of use and inherent scalability, a recent insight from *Gal & Ghahramani*^31,32,35^is particularly promising for use in medical settings. Using dropout networks^36,37^, where subsets of units are inactivated during training to avoid overfitting, one can compute an approximation to the posterior distribution by sampling multiple predictions with dropout turned on. This allows one to perform approximate but efficient Bayesian inference by using existing software implementations in a straightforward way. Another advantage of this approach is that it can be applied to already trained networks.

Here we assess whether this allows us to retrieve informative uncertainty estimates for a large-scale, real world disease detection problem and contrast it against straightforward alternatives: i) the conventional network output via *standard dropout* and ii) Gaussian processes^38^ (GPs). Diabetic retinopathy (DR) is one of the leading causes of blindness in the working-age population of the developed world^39^. If the symptoms are detected in time, progress to vision impairment can be averted but the existing infrastructure is insufficient and manual detection is time-consuming. With the increase in the global incidence of diabetes^40^, clinicians now recognize the need for a cost-effective, accurate and easily performed automated detection of DR to aid the screening process^14,39,41–43^. Previous recommendations of the British Diabetic Association (now Diabetes UK) are often cited as 80% sensitivity and 95% specificity [42,44,45, and references therein] but the current minimum thresholds set by the NHS Diabetes Eye Screening programme are 85% sensitivity and 80% specificity for sight-threatening diabetic retinopathy^16^.

Using a Bayesian DNN, we achieve state-of-the-art results for diabetic retinopathy detection on the publicly available dataset Messidor^46^. The computed measure of uncertainty allowed us to refer a subset of difficult cases for further inspection, resulting in substantial improvements in detection performance in the remaining data. While it is sometimes believed that the conventional network output captures the networks uncertainty, neither this nor a GP alternative were competitive under the decision referral scenario. This finding generalized across different model architectures, detection tasks and datasets. In practice, patients whose samples result in uncertain decisions would either be sent for further screening tests or referred directly to a specialist. We further explore the causes of uncertainty in our scenario. Intuitions illustrated on a 2D toy problem are used to understand how uncertainty might behave in the high-dimensional image space. This allowed us to predict the kind of application relevant scenarios for which the assessed uncertainty is informative.

## Results

Here we tackle two major questions: first, we evaluate whether model uncertainty obtained from deep disease detection networks at test time is useful for ranking test data by their prediction performance without knowing the latter. In the second part, we open the black box in order to learn what renders predictions uncertain.

### Predicting diabetic retinopathy with a measure of (un)certainty

#### Diabetic retinopathy datasets

We used a DNN-based approach to detect diabetic retinopathy (DR) from fundus images. Our main dataset used for training is taken from a previous Kaggle competition^47^. This dataset consists of 35,126 training images and 53,576 test images, graded into five stages of DR by clinicians according to the following scale^48^: 0 - No DR, 1 - Mild, 2 - Moderate, 3 - Severe and 4 - Proliferative DR. The percentage of images labelled with No DR is about 73% in both the training and test dataset.

In order to measure the true generalization of our insights we in addition applied all networks to the publicly available Messidor dataset^46^. This dataset comprises 1,200 images divided into the following categories: 0 - No DR, 1 - Mild non-proliferative, 2 - Severe non-proliferative, 3 - Most serious DR.

#### Disease detection tasks

Because the question of whether a patient has to be sent to a physician at all, is of high priority, we reduced the problem to a binary classification task. Therefore we split the data into a “healthy” versus “diseased” set by grouping some of the classes. In order to analyse how model uncertainty behaves for different tasks, we varied the disease onset level. If set to 1, the classes except for 0 are in the “diseased” category resulting in a detector for mild DR (or more severe) whereas for disease onset level 2, classes {0,1} are considered “healthy” and moderate DR (or more severe levels) are in the “diseased” group.

#### Network architectures

We used two different network architectures for our experiments: (1) Two networks trained for the questions at hand. (2) The publicly available network architecture and weights^49^ provided by the participant who scored very well in the Kaggle DR competition^47^, which we will call *JFnet*.

The JFnet comprises 13 convolutional layers, 3 fully connected layers and a concatenation layer combining information from the contralateral eyes of a patient. Convolutional layers are interleaved with 5 max pooling layers, fully connected layers are interleaved with two feature pooling and dropout (*p*_*drop*_ = 0.5) layers each. All non-linearities are ReLUs^50^ or Leaky ReLUs^51^ (leakiness 0.5) except for the softmax output layer^52^. We recast the original model’s five output units (trained for Kaggle DR’s level discrimination task) to our binary tasks by summing the output of respective units.

Our own network architecture was inspired by the monocular part of the JFnet (which in turn is VGG-like^53^, a standard CNN architecture that has been shown to perform well in a variety of applications), with the fully connected part replaced by the concatenation of a global mean and a global max pooling layer, followed by a softmax output layer. In contrast to the JFnet, our networks do not rely on both images of a given patient being present, i.e. they do not perform eye blending. Furthermore, they have more network parameters that are treated in a Bayesian manner^32^, because we added dropout (*p*_*drop*_ = 0.2) after each convolutional layer for which reason we denote these networks as *Bayesian* convolutional neural networks (*BCNN*s).

#### Bayesian model uncertainty

For the main part of the paper, we measured the uncertainty associated with the predictions of the DNNs described above, exploiting a relationship between dropout networks and a Bayesian posterior^31^. For evaluation purposes, we compare against alternative uncertainty measures where appropriate. Typically, the softmax output of a classification network denotes a single prediction given a sample. In case of DR detection from a fundus image (see fig. 1a left, for a “diseased” example) a trained network would output the probability that the given image is “diseased” (fig. 1a, right). The softmax probability is based on a single set of network parameters, whereas in a Bayesian setting one aims for the predictive posterior (compare (Eq. 2), i. e. a distribution over predictions (in our case the softmax values) obtained by integrating over the distribution over possible parameters.

The predictive posterior of a neural network is hard to obtain. However, Gal and colleagues^31^ showed that by leaving dropout turned on at test time, we can draw Monte Carlo samples from the approximate predictive posterior (for details see Methods). We will summarize each predictive posterior distribution by its first two moments. The predictive mean µ_*pred*_ (eq. 7) will be used for predictions and the predictive standard deviation σ_*pred*_ (eq. 8) as the associated uncertainty.

**Figure. 1.**
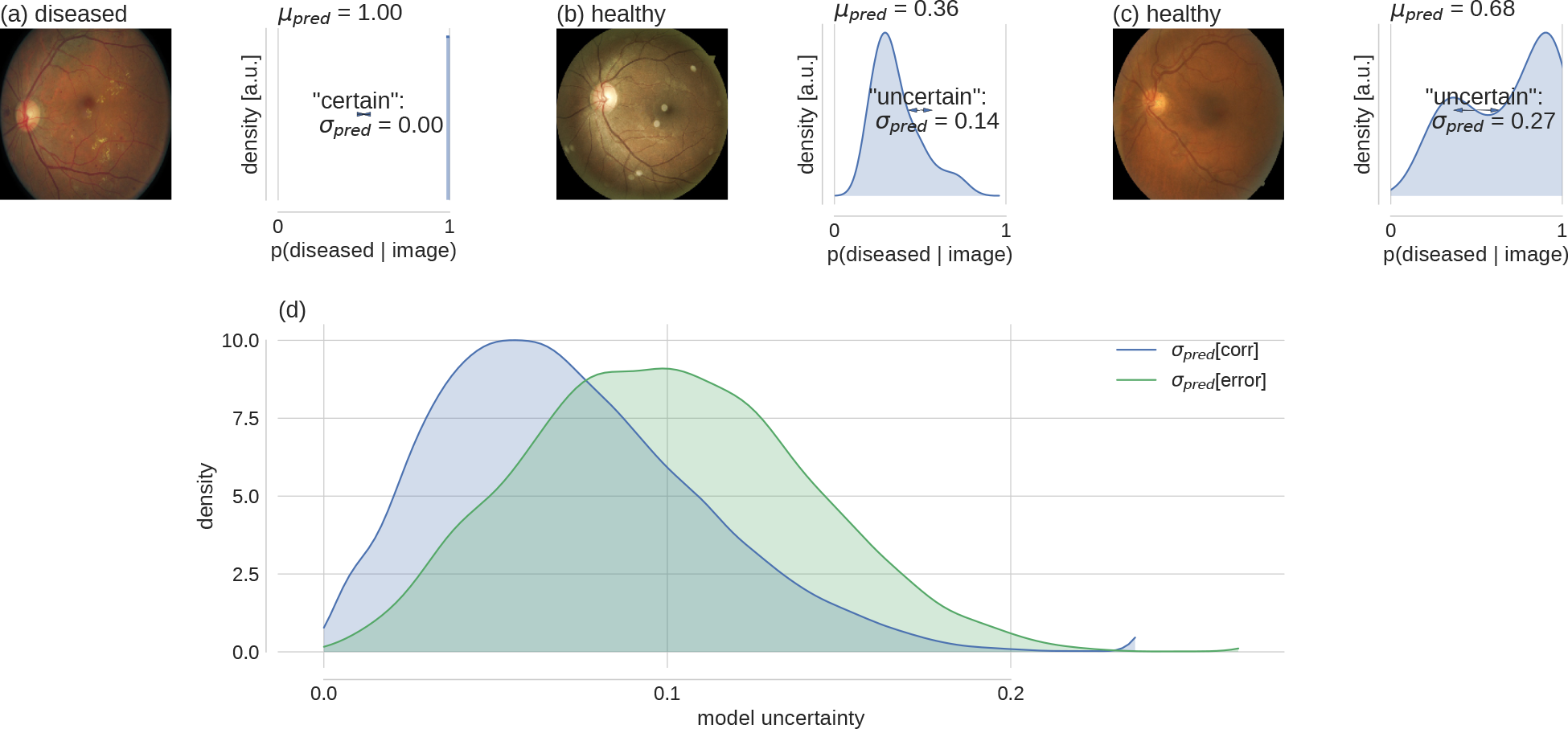
Bayesian model uncertainty for diabetic retinopathy detection. **(a)-(c), left:** Exemplary fundus images with human label assignments in the titles. (**a**)-(**c**), **right**: Corresponding approximate predictive posteriors (Eq. 6) over the softmax output values p(diseased|image) (Eq. 1). Predictions are based on µ_*pred*_ (Eq. 7) and uncertainty is quantified by σ_*pred*_ (Eq. 8). Examples are ordered by increasing uncertainty from left to right. (**d**) Distribution of uncertainty values for all Kaggle DR test images, grouped by correct and erroneous predictions. Label assignment for “diseased” was based on thresholding µ_*pred*_ at 0.5. Given a prediction is incorrect, there is a strong likelihood that the prediction uncertainty is also high.

Based on a fundus image, a DNN can be certain (fig. 1a) or more or less uncertain (fig. 1b–c) about its decision, as indicated by the width of the predictive posterior distribution: For example, an image can be classified as certainly diseased, where all sampled predictions are 1.0, such that σ_*pred*_ = 0 (fig. 1a). A different example is classified as “healthy”, but the network predictions are more spread out (σ_*pred*_ = 0.14) (fig. 1b). The predicted label is still correct, because µ_*pred*_ = 0.36 < 0.5. Finally, some examples produce high uncertainty in the DNN (σ_*pred*_ = 0.27) and result in an erroneous “diseased” prediction (σ_*pred*_ = 0.68 > 0.5) (fig. 1c).

If high model uncertainty was indicative of erroneous predictions, this information could be leveraged to increase the performance of the automated system by selecting appropriate subsets for further inspection. Indeed, model uncertainty was higher for incorrect predictions (fig. 1d). This means that σ_*pred*_ (a quantity that can be evaluated at test time) can be used to rank order prediction performance (a quantity unknown at test time), in order to mimic the human clinical work flow. In face of uncertain decisions, further information should be obtained.

Importantly, model uncertainty as quantified by σ_*pred*_ adds complementary information to the conventional network output as quantified by *p*(*diseased\image*) (eq. 1), i.e. with dropout turned off at test time. Specific softmax values do not determine the precise values that σ_*pred*_ assumes (fig. 2). This decouples prediction uncertainty as measured by *p(diseased\image)* from model uncertainty. Lower probabilities that an image is diseased are associated with a larger range of uncertainties while high probabilities that an image is diseased are confined to smaller uncertainties, indicating that if an image can be classified as diseased, this typically happens with confidence. In contrast, healthy is a much less crisp concept, where variation among individuals can lead to significant uncertainty in judgement.

**Figure. 2.**
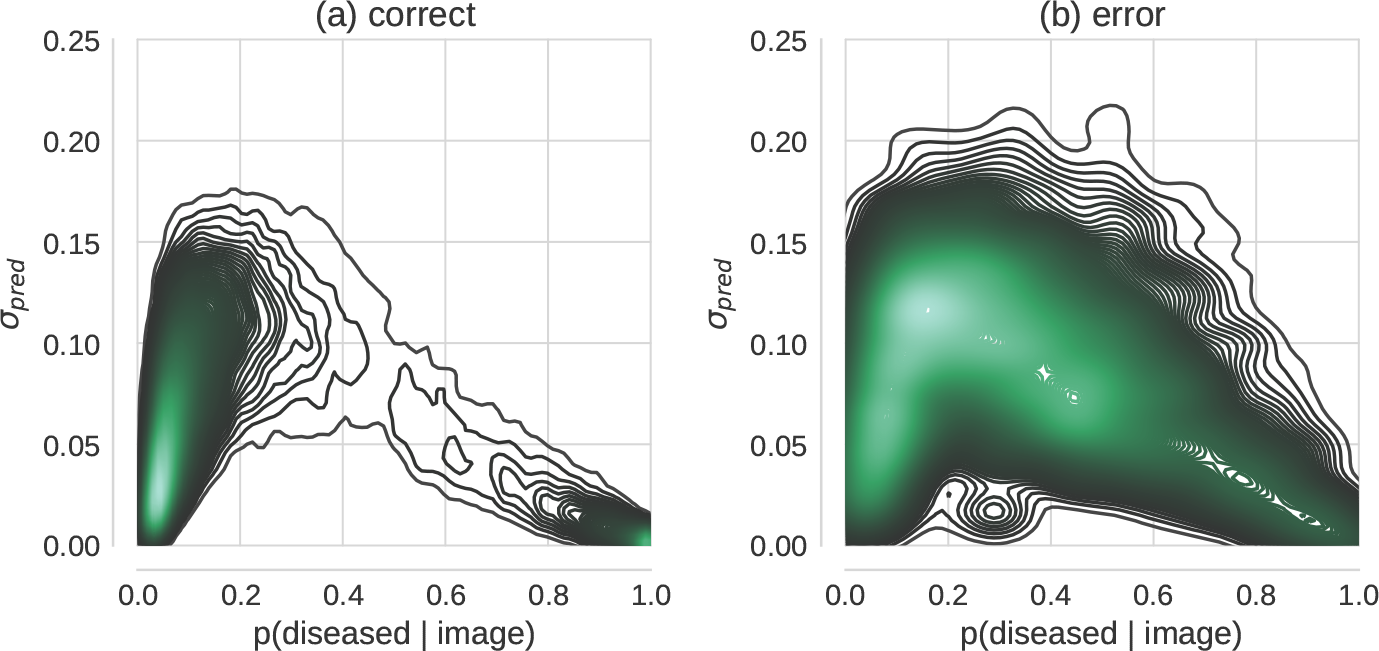
Relation between Bayesian model uncertainty σ_*pred*_ and maximum-likelihood, i.e. conventional softmax probabilities*p*(*diseased\image*). Each subplot shows the 2-dimensional density over Kaggle DR test set predictions conditioned on: correctly (**a**) vs. erroneously (**b**) classified images respectively.

### Uncertainty rank orders prediction performance

#### Performance improvement via uncertainty-informed decision referral

To test whether we could exploit the uncertainty measurement proposed above to mimic the clinical workflow and refer patients with uncertain diagnosis for further testing, we performed predictions (using the BCNN trained for disease onset 1 on the Kaggle DR training images) for all Kaggle DR test images and sorted the predictions by their associated uncertainty. We then referred predictions based on various levels of tolerated uncertainty for further diagnosis and measured the accuracy of the predictions (threshold at 0.5) for the remaining cases (fig. 3a).

**Figure. 3.**
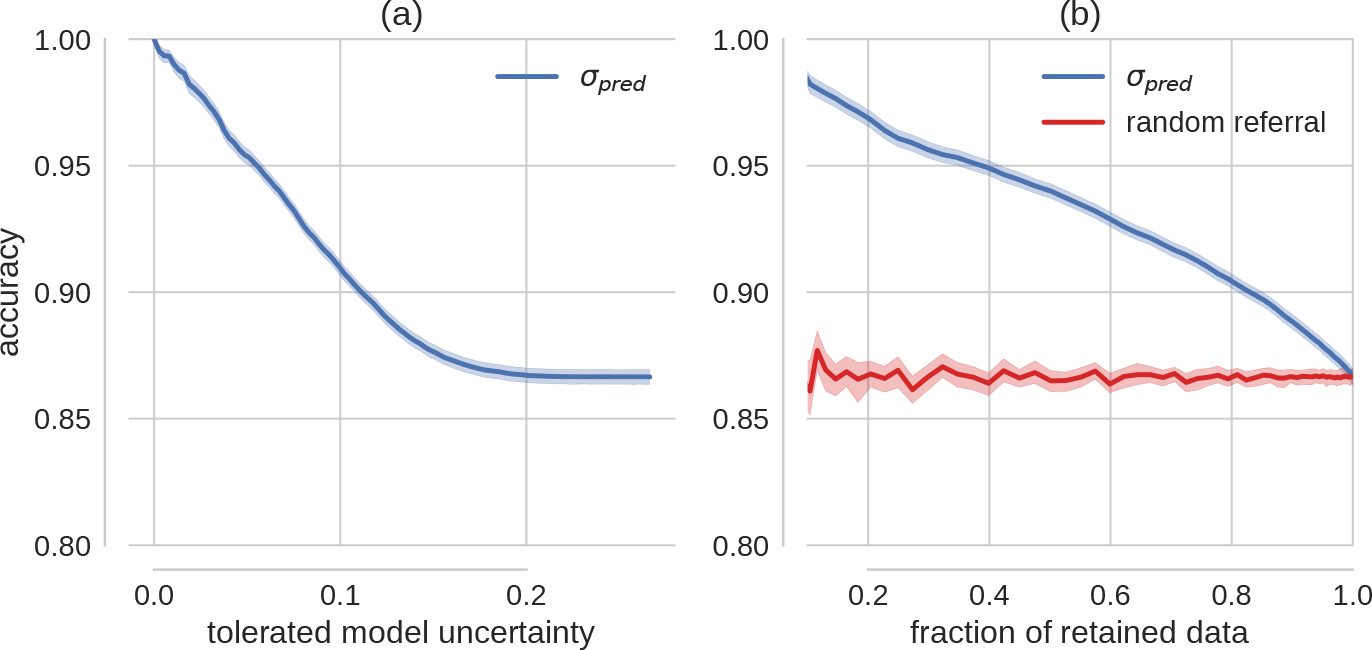
Improvement in accuracy via uncertainty-informed decision referral. (**a**) The prediction accuracy as a function of the tolerated amount of model uncertainty. (**b**) Accuracy is plotted over the retained data set size (test data set size - referral data set size). The red curve shows the effect of rejecting the same number of samples randomly, that is without taking into account information about uncertainty. Exemplarily, if 20% of the data would be referred for further inspection, 80% of the data would be retained for automatic diagnostics. This results in a better test performance (accuracy ≥ 90%, point on blue curve) on the retained data than on 80% of the test data sampled uniformly (accuracy ≈ 87%, point on red curve). Uncertainty informed decision referral derived from the conventional softmax output cannot achieve consistent performance improvements (fig. 4).

We observed a monotonic increase in prediction accuracy for decreasing levels of tolerated model uncertainty, which translates to the same behaviour when monitoring the fraction of retained, that is automatically diagnosed, data instead (fig. 3b, blue curve). As a control experiment, we compared with randomly selected data samples, that is without using uncertainty information (fig. 3b, red curve). For less than 2% decisions referred for further inspections, the 95% confidence intervals of the two scenarios are already non-overlapping. Uncertainty is hence informative about prediction performance, here measured by accuracy.

#### Performance improvement for different costs, networks, tasks and datasets

Here we build on the idea of uncertainty informed decision referral introduced above (fig. 3) and assess whether performance improvements are robust across different settings. So far (fig. 1–3), predictions had been converted to labels by thresholding the network output at 0.5. In a medical setting however, different costs are associated with false positive and false negative errors.

These can be controlled by the decision threshold at which the diseased probability given an image is converted to the category “diseased”. A complete picture can be obtained by the decision system’s receiver-operating-characteristic, which monitors *sensitivity* over *1 - specificity* pairs for all conceivable decision thresholds. The quality of such a ROC curve can be summarized by its area under the curve (AUC), which varies between 0.5 (chance level) and 1.0 (best possible value). Importantly, ROC characteristics allow us to assess model uncertainty independent of prediction uncertainty.

ROC AUC improves monotonically with decreasing levels of tolerated uncertainty (fig. 4a, left, blue curve). In addition, ROC curves for all Kaggle test images as well as under 10,20 and 30% decision referral reveal that both sensitivity and specificity consistently improved (fig. 4a, right). These results were found to be robust for a variety of settings, that is for our different networks/tasks (disease onset 1 (fig. 4, left double column) vs. disease onset 2 (fig. 4, right double column)) as well as when applied to the completely independent Messidor database (fig. 4, 2nd row). For comparison with a different network architecture we refer to Supplementary figure S1.

**Figure. 4.**
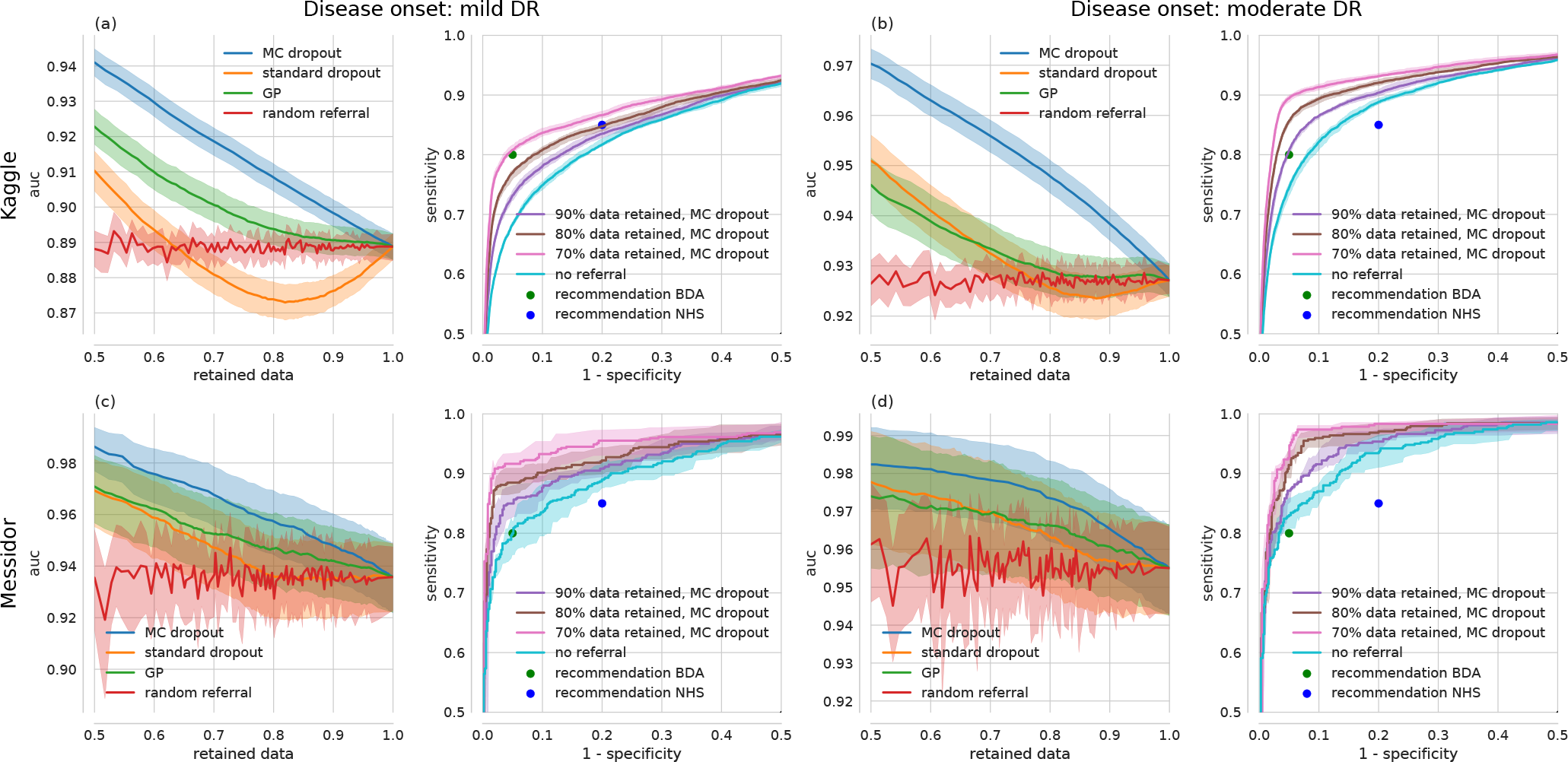
Improvement in receiver-operating-characteristics via uncertainty-informed decision referral for different networks/tasks (left vs. right double column), datasets (1st vs. 2nd row) and methods (1st and 3rd single column). (**a, left**) ROC AUC over the fraction of retained data under uncertainty informed (MC dropout: blue, Gaussian process: green, standard dropout: orange) and random (red) decision referral for a Bayesian CNN, trained for disease onset 1 and tested on Kaggle DR. (**a, right**) Exemplary ROC curves under decision referral using the best method from (a, left), that is MC dropout. ROC curves improve when increasing the number of referred samples (90/80/70% retained data: purple/brown/pink curves respectively) as compared to no referral (turquoise). Panels (b)-(d) have the same layout. National UK standards for the detection of sight-threatening diabetic retinopathy (in^55^ defined as moderate DR) from the BDA (80%/95% sensitivity/specificity, green dot) and the NHS (85%/80% sensitivity/specificity, blue dot) are given in all subpanels with ROC curves. (**b**) same as (a), but for disease onset 2. (**c**) Same network/task as in (a), but tested on Messidor. (**d**) Same network/task as in (b), but tested on Messidor.

The application to Messidor provides a report of the true generalization performance, because it had never been used for either of our networks. Our networks are thus robust against the distributional shift between Kaggle and Messidor which will be analysed further below in more detail.

Even though our primary aim was not to achieve high performance, we surpassed the requirements of both the NHS and the British Diabetic Association (fig. 4, blue and green dots respectively) for (automated) patient referral for several settings and perform on par with the non-ensembling approach of *Antal & Hajdu*^44^. We also performed similar ensembling^18,44^, by selecting an optimal (forward-backward search while monitoring AUC) ensemble of 100 networks from a much larger pool of dropout networks by controlling the random seeds. Performance improvements however were marginal and did not generalize to test data, probably because this compromises the stochastic nature of the regularizing effects of dropout. For a summary of the different configurations and comparison with the state-of-the-art we refer to Table 1.

**Table 1.**
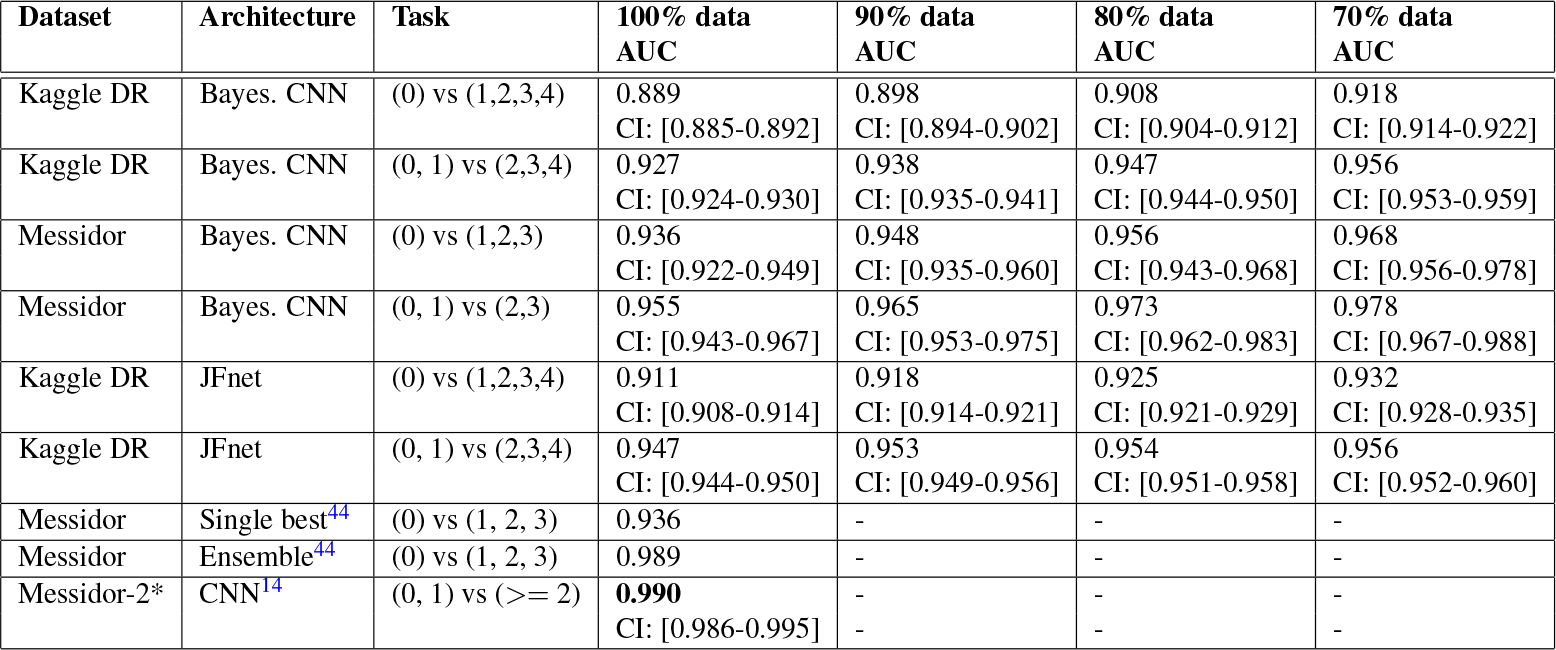
Model performance (measured by AUC) with two different datasets, architectures and tasks when data with higher uncertainty levels is referred to further inspection. *For Messidor-2^46,56^no labels are publicly available for comparison with our networks.

| Dataset | Architecture | Task | 100% data<br>AUC | 90% data<br>AUC | 80% data<br>AUC | 70% data<br>AUC |
| --- | --- | --- | --- | --- | --- | --- |
| Kaggle DR | Bayes. CNN | (0) vs (1,2,3,4) | 0.889<br>CI: [0.885-0.892] | 0.898<br>CI: [0.894-0.902] | 0.908<br>CI: [0.904-0.912] | 0.918<br>CI: [0.914-0.922] |
| Kaggle DR | Bayes. CNN | (0, 1) vs (2,3,4) | 0.927<br>CI: [0.924-0.930] | 0.938<br>CI: [0.935-0.941] | 0.947<br>CI: [0.944-0.950] | 0.956<br>CI: [0.953-0.959] |
| Messidor | Bayes. CNN | (0) vs (1,2,3) | 0.936<br>CI: [0.922-0.949] | 0.948<br>CI: [0.935-0.960] | 0.956<br>CI: [0.943-0.968] | 0.968<br>CI: [0.956-0.978] |
| Messidor | Bayes. CNN | (0, 1) vs (2,3) | 0.955<br>CI: [0.943-0.967] | 0.965<br>CI: [0.953-0.975] | 0.973<br>CI: [0.962-0.983] | 0.978<br>CI: [0.967-0.988] |
| Kaggle DR | JFnet | (0) vs (1,2,3,4) | 0.911<br>CI: [0.908-0.914] | 0.918<br>CI: [0.914-0.921] | 0.925<br>CI: [0.921-0.929] | 0.932<br>CI: [0.928-0.935] |
| Kaggle DR | JFnet | (0, 1) vs (2,3,4) | 0.947<br>CI: [0.944-0.950] | 0.953<br>CI: [0.949-0.956] | 0.954<br>CI: [0.951-0.958] | 0.956<br>CI: [0.952-0.960] |
| Messidor | Single best<sup>44</sup> | (0) vs (1, 2, 3) | 0.936 | - | - | - |
| Messidor | Ensemble<sup>44</sup> | (0) vs (1, 2, 3) | 0.989 | - | - | - |
| Messidor-2* | CNN<sup>14</sup> | (0, 1) vs ($\geq 2$) | **0.990**<br>CI: [0.986-0.995] | - | - | - |

The better performance for moderate DR detection (onset 2) as compared to mild DR detection (onset 1) across networks and datasets is in line with the more pronounced expression of symptoms as the disease progresses. Comparison across datasets reveals that for both tasks, the models performed better on Messidor than on Kaggle data (compare fig. 4a vs. c and b vs. d). Specifically, we achieved both the BDA and NHS requirements on Messidor without having to refer decisions whereas for Kaggle data we have to refer 0 - 30% of the data, depending on the recommendation, task difficulty and network capacity. It has been reported previously that about 10%^54^ of the Kaggle DR images were considered ungradable according to national UK standards. We want to emphasize that the proposed uncertainty informed decision referral did not rely on labels for such cases, that is we could detect performance impeding images without training a separate, supervised detection algorithm. To what extent images associated with low model confidence relate to clinically relevant cases or ungradable images will be analysed in the section about *what causes uncertainty*.

#### Comparison with alternative uncertainty measures

We rendered large-scale disease detection networks Bayesian by using the *MC dropout* approach put forward by Gal and colleagues^60^ because it is theoretically sound, easy to implement and computationally efficient. It remains however an open question how MC dropout compares to alternative uncertainty measures. The decision referral scenario can serve as a minimal benchmark for comparing uncertainty methods. Here we contrast against two straightforward and seemingly appealing alternatives, that is the conventional network output (*standard dropout*) and *GPs*^38^ fit to the penultimate layer features.

Because the conventional network output denotes the probability that a given image is diseased, the decision referral scenario can reveal whether a Bayesian approach is indeed necessary. Because we do not have a distribution over predictions in case of standard dropout, σ_*pred*_ is undefined. We can however resort to quantifying the uncertainty about a prediction by the binary entropy *H*(*p*) = −(*p*log*p*+(1 – *p*) log (1 − *p*)) instead. Entropy is theoretically grounded and applicable to the Bayesian and conventional network outputs as well as the GP outputs. For our Bayesian networks, the entropy performs comparable to σ_*pred*_ as shown in Supplementary Figure 2 and can hence be used as a drop-in replacement for the decision referral scenario. In case of using the uncertainty derived from standard dropout network (orange curves in fig. 4), the performance improvement under decision referral is lower than with MC dropout (blue curves in fig. 4). When referring up to ≈ 30% of the decisions considered most uncertain in case of detecting mild DR, standard dropout performs even worse than random referral (fig. 4, left, orange vs. red curves). This means that the class probabilities obtained via standard dropout are miscalibrated (see as well fig. 2) and not suited for performance improvements under decision referral. Because we need to compute either the mean (for *H*(*µ*_*pred*_)) or the standard deviation σ_*pred*_ of the (approximated) predictive posterior, we conclude that we do need Bayesian methods to achieve our results.

Given this, GPs could in principle constitute an alternative approach to MC dropout (see Supplementary Information). While they may theoretically seem more appealing than dropout-uncertainty, they scale badly with both the dimensionality of the feature space and the size of the dataset. The standard GP learning exhibits a runtime complexity of *O*(*N*^3^) and memory complexity of *O*(*N*^2^), where *N* is the size of the dataset^38,74^. To render the application of GPs feasible for disease detection from a large collection of high-dimensional medical images, we adopted the minibatch approach and the 512-dim activations of the last hidden layers of our Bayesian networks were used as input to the GP classifiers that are equipped with neural network covariance functions (see Supplementary notes). As it was necessary to work with a single feature vector per image, conventional dropout was used to obtain them. While this was sufficient to achieve similar performance without referring data (blue and green curves overlap strongly for 100% retained data in the 1st and 3rd column of fig. 4), such a greedy training of GPs, which are essentially shallow neural networks, on the last hidden layer activations of a deep network inherently suffers from two issues: i) the loss of the Bayesian treatment of MC dropout, and ii) the lack of access to the full stack of knowledge from earlier layers. As a result, GPs cannot keep up with our Bayesian networks under any decision referral scenario (fig. 4, green vs. blue curves for both datasets and disease detection tasks).

In addition to architectural disadvantages, we report on practical difficulties of using GPs. Despite the promises of Expectation Propagation (EP) for approximate inference for GP classification (see Supplementary notes), we could not utilize it in our experiments. More specifically, the inference routine failed to converge during our trials, and we had to resort to a simpler approximate inference algorithm, namely Laplace Approximation (LA). Given that this method has been reported to substantially underestimate the mean and covariance of a GP, especially in high-dimensional spaces (see Supplementary notes), we speculate that it prevented the uncertainty estimates from being as good as they could be with EP, and ultimately hindered the performance of GPs under the decision referral scenario.

As the Bayesian network approach performed best throughout, the following sections will exclusively analyse the properties of the uncertainty derived from MC dropout. Some aspects of what causes uncertainty are anyway fundamentally determined by the fact of dealing with a classification problem and should qualitatively hold for different uncertainty measures.

### What causes uncertainty?

Next we asked what causes the networks to consider the prediction about an image uncertain. In order to build an intuitive understanding of uncertainty estimates, we trained a simple Bayesian neural network (3 hidden layers with 100 units each) with dropout layers interleaved (*p*_*drop*_ = 0.5) on a 2D toy classification problem (fig. 5).

**Figure. 5.**
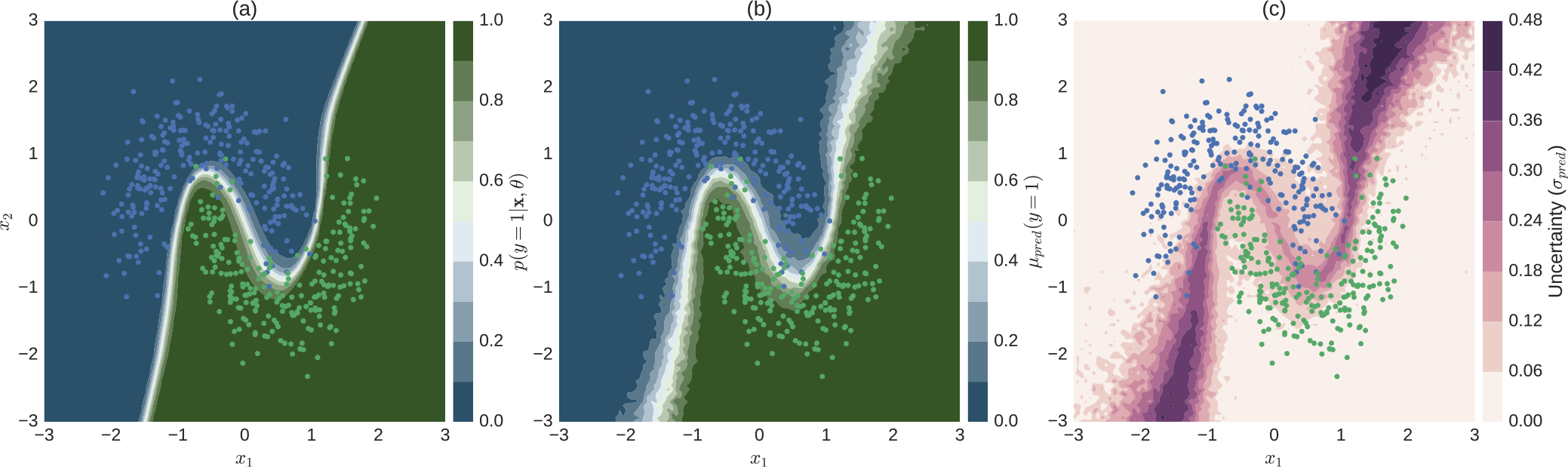
Illustration of uncertainty for a 2D binary classification problem. (**a**) Conventional softmax output obtained by turning off dropout at test time (eq. 1). (**b**) Predictive mean of approximate predictive posterior (eq. 7). (**c**) Uncertainty, measured by predictive standard deviation of approximate predictive posterior (eq. 8). The softmax output (a) is overly confident (only a narrow region in input space assumes values other than 0 or 1) when compared to the Bayesian approach (b, c). Uncertainty (c) tends to be high for regions in input space through which alternative separating hyperplanes could pass. Colour-coded dots in all subplots correspond to test data the network has not seen during training.

The network learns the non-linear hyperplane (defined by *p*(*y* = 1|**x, *θ***) = 0.5) that separates the two classes (fig. 5a) shown as the network's softmax output when evaluated traditionally, that is with dropout turned off at test time. The first (fig. 5b, eq. 7) and second moment (fig. 5c, eq. 8) of the approximate predictive posterior (eq. 6) in turn are more spread out along directions orthogonal to the separating hyperplane given by the conventional softmax output. This is because the Bayesian perspective models a distribution over possible separating hyperplanes. In contrast to the conventional network output, µ_*pred*_ and σ_*pred*_ are more related. Incidentally, this was also true in the high-dimensional real world setting, where mean and the standard deviation of the predictive posterior resulted in similar uncertainty judgements (fig. S2). Note that regions in the input space that have high probabilities of belonging to either class in the non-Bayesian setting (fig. 5a) may still carry substantial uncertainties in the Bayesian setting (fig. 5c), which is in line with the high-dimensional case as illustrated by Figure (2).

In order to evaluate the relationship of uncertainty with respect to the class boundary, we devised an experiment that makes use of the gradual progression of disease levels from 0 to 4 as provided by physicians in case of Kaggle DR data. We probed what happened to images with different “distances” from the healthy/diseased boundary defined by the disease onset level of a given task. To this end, we quantified the proportion of the different disease levels in the data referred for further inspection for various levels of tolerated uncertainty (fig. 6).

**Figure. 6.**
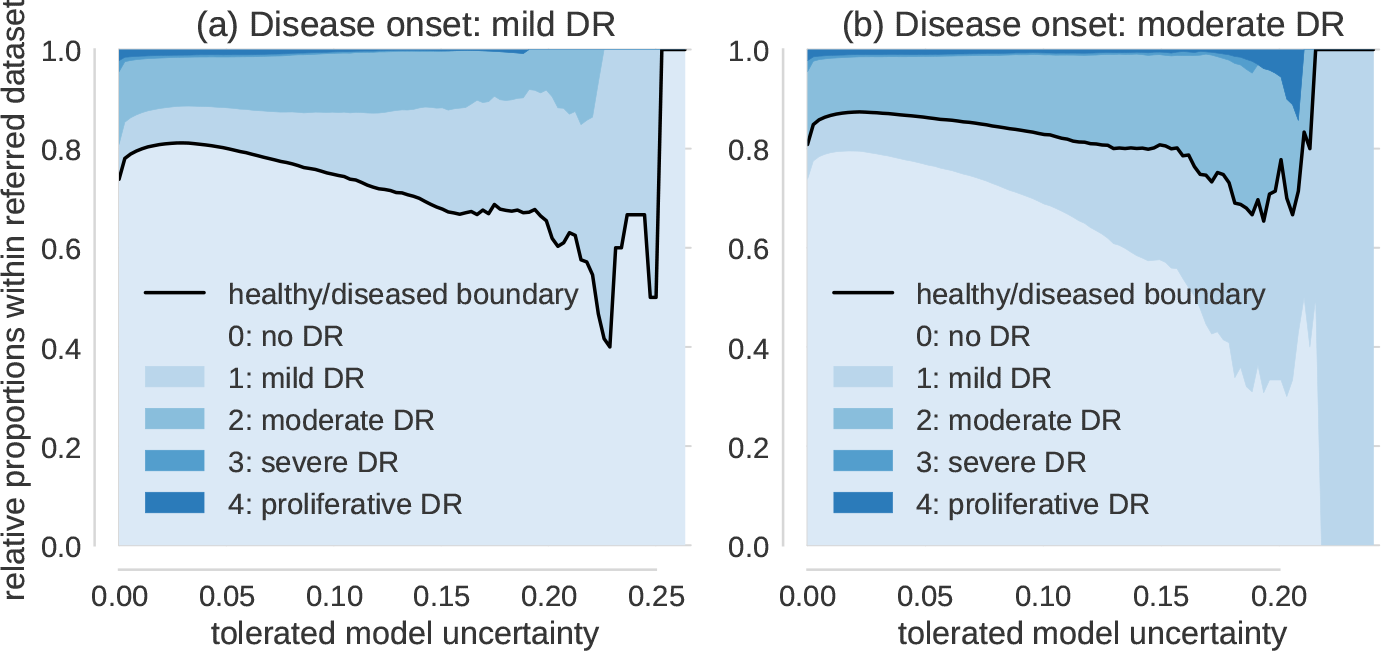
Proportion of disease levels in referred datasets. The value on the x-axis indicates the uncertainty of a sample to be tolerated for automatic diagnosis. All samples in the referral dataset have thus uncertainties of at least the value on the x-axis. The relative proportion of disease levels for tolerated uncertainty = 0 corresponds to the prior. (**a**) Disease onset level is mild DR (1). Disease levels 0 and 1 neighbour the healthy/diseased boundary (black) and dominate the referral dataset for high but not intermediate uncertainty. (**b**) Disease onset level is moderate DR (2). In analogy to (a), disease levels 1 and 2 neighbour the healthy/diseased boundary (black) and dominate the decision referral populations with high in contrast to intermediate uncertainties.

If no model uncertainty is tolerated, we observe the prior distribution (shown on the vertical axis at σ_*pred*_ = 0) of disease levels because all data is referred. If instead only the most uncertain cases are referred, the contribution of those disease levels that are adjacent to the healthy/diseased boundary (black lines in fig. 6a & b) is increased. For mild DR defining the disease onset and large tolerated uncertainties, disease levels 0 and 1 dominate the pool of referred data (fig. 6a). If we shifted the disease onset to moderate DR, in an analogous manner disease levels 1 and 2 dominate the referred data sets for high uncertainties (fig. 6b). In an intermediate regime however, such as e.g. around an uncertainty of 0.1 for which we still refer less than 25% of the data, the relative contribution of disease levels is already resembling the prior. Taken together with the fact that we improve performance throughout at least up to 50% data referral (compare fig. 3&4) it is not only those samples that neighbour the class boundaries that carry meaningful uncertainties as determined by the networks.

As a side note, depending on the therapeutic possibilities - moderate DR detection (fig. 6a) might be preferable to mild DR detection (fig. 6a) as the uncertainty still detected level 1 patients in the latter case but reduced the amount of healthy patients sent for referral.

In the following we devised an experiment that aimed to evaluate the algorithm's referral decisions against physicians’ decisions. For that purpose we made use of the availability of both eyes’ images for each patient in case of Kaggle DR data. Even though therapeutic intervention is typically based on a patient level diagnosis, the contra-lateral eye of a given patient may be in a different state and therefore carry a different label. A strong correlation of the two eyes’ disease states was leveraged to improve performance by many Kaggle competition participants^47^. However, even after compilation of the 5-class problem to the binary disease detection problem, 5 - 10% of images categorized as diseased have images from the contra-lateral eye with a disagreeing label.

Whether the corresponding patients are diseased or not is therefore undefined and they should be subject to referral. By measuring the proportion of images whose contra-lateral ground truth label is different for the referred and retained data sets respectively (fig. 7), we could analyse to what extent the model uncertainty reflects a physician’s uncertainty. Exemplarily (fig. 7a), if the tolerated model uncertainty were 0.2, only ≈ 8% of the retained images (σ_*pred*_ < = 0.2) belong to a patient with ambiguous disease state whereas nearly 20% of the referred images (σ_*pred*_ > 0.2) belong to a patient with ambiguous disease state. Throughout, images from patients with one healthy and one diseased eye are more likely to be referred for further inspection than retained (fig. 7). For both disease detection tasks (fig. 7 a/b for mild/moderate DR as disease onset respectively) this is particularly pronounced in the regime of high uncertainty.

**Figure. 7.**
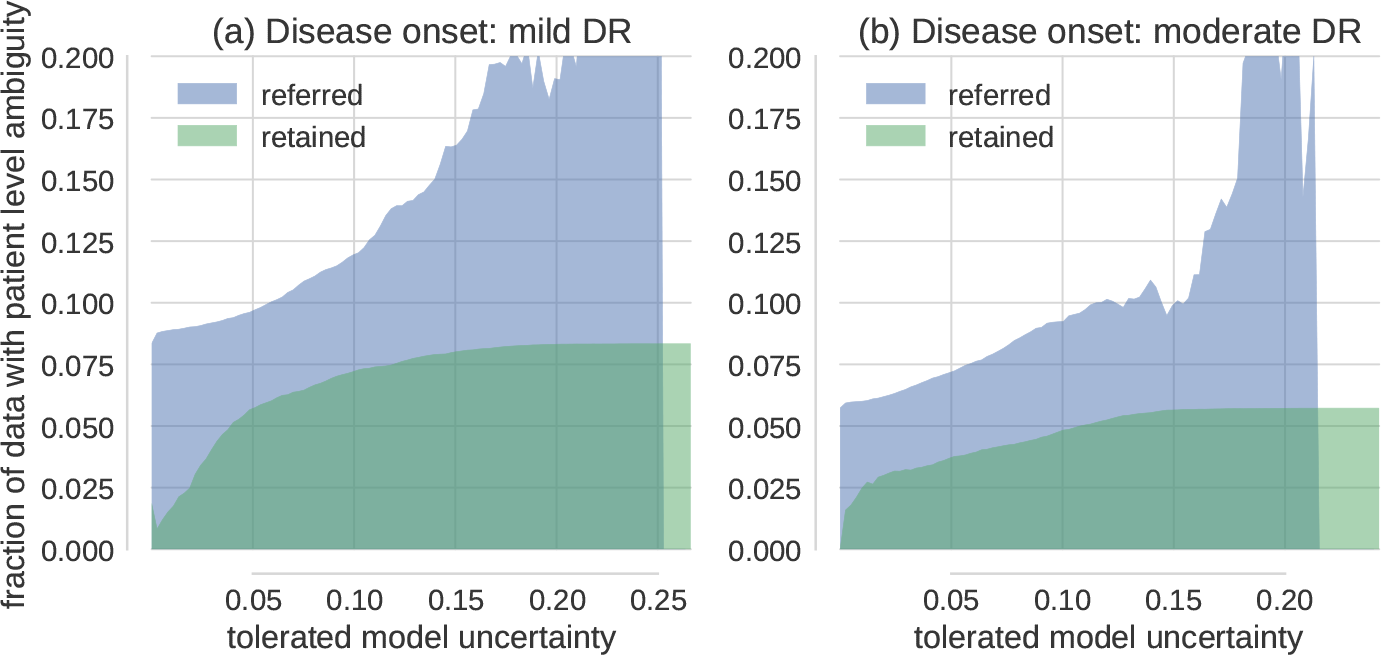
Decision referral of images from ambiguous patients. (**a**) Disease onset is mild DR (1). (**b**) Disease onset is moderate DR (2). Both subplots show the relative proportion of images from ambiguous patients in the referred (blue) and retained (green) data buckets for various tolerated uncertainty values. Patient level ambiguity is defined by images whose contra-lateral eye (from the same patient) carries a different label. Note that the decision referral of images is based on the uncertainty from a single image. Ground truth labels and the contra-lateral eye information are only used as meta information for evaluation purposes. Especially in the high uncertainty regime, images from ambiguous patients are more likely to be referred for further inspection than accepted for automatic decision. This is in line with how a physician would decide because ambiguous patients have an undefined disease state and should be subject to further examination.

### Uncertainty about unfamiliar data samples

Ideally, model uncertainty about the predicted class would be high for data not trained to be recognized or even be sensitive to adversarial attacks^57^. In this case, one could use uncertainty not only to determine which images are hard to diagnose and require further inspection, but also to sort out unusable data. Unfortunately, uncertainty about a discriminative model (family) is not necessarily suited to detect samples “far” from the training data. In two dimensions (compare fig. 5c) it is easy to see that regions that are both far away from the data and carry high model uncertainty are not isotropically distributed with regard to the data but rather task dependent. With increasing dimensionality of the input space, more separating hyperplanes are conceivable to solve a given task, attributing non-zero uncertainties to a larger fraction of the input space. Nevertheless, the task dependency is built into the model. In the following we show for the high-dimensional scenario that the task and dataset difficulty influence the separability of unfamiliar data samples from the distribution the network was trained for.

The space of images with content unknown to the disease detection networks was sampled by performing predictions with associated uncertainties on the 2012 Imagenet validation set (49101 coloured, non-fundus images from 1000 different categories)^58^ (fig. 8 a,b). Interestingly, Messidor images have the lowest average uncertainty, followed by Kaggle and Imagenet samples, especially for disease onset 2. The networks perform much better on the Messidor dataset than on the Kaggle dataset, and disease detection for onset 2 is much easier than for onset 1, indicating that in relatively easy tasks/datasets uncertainty does to some extent serve to detect out-of-sample images. If the distinction between “healthy” and “diseased” was clear cut such as in the case of many classes used for classical computer vision tasks, it is well conceivable that the uncertainty distributions would be well separated for the known and unknown classes^18^. Because we observe substantial uncertainty about the presence of DR, the detection of unfamiliar data samples is however obscured, at least for the Kaggle dataset.

**Figure. 8.**
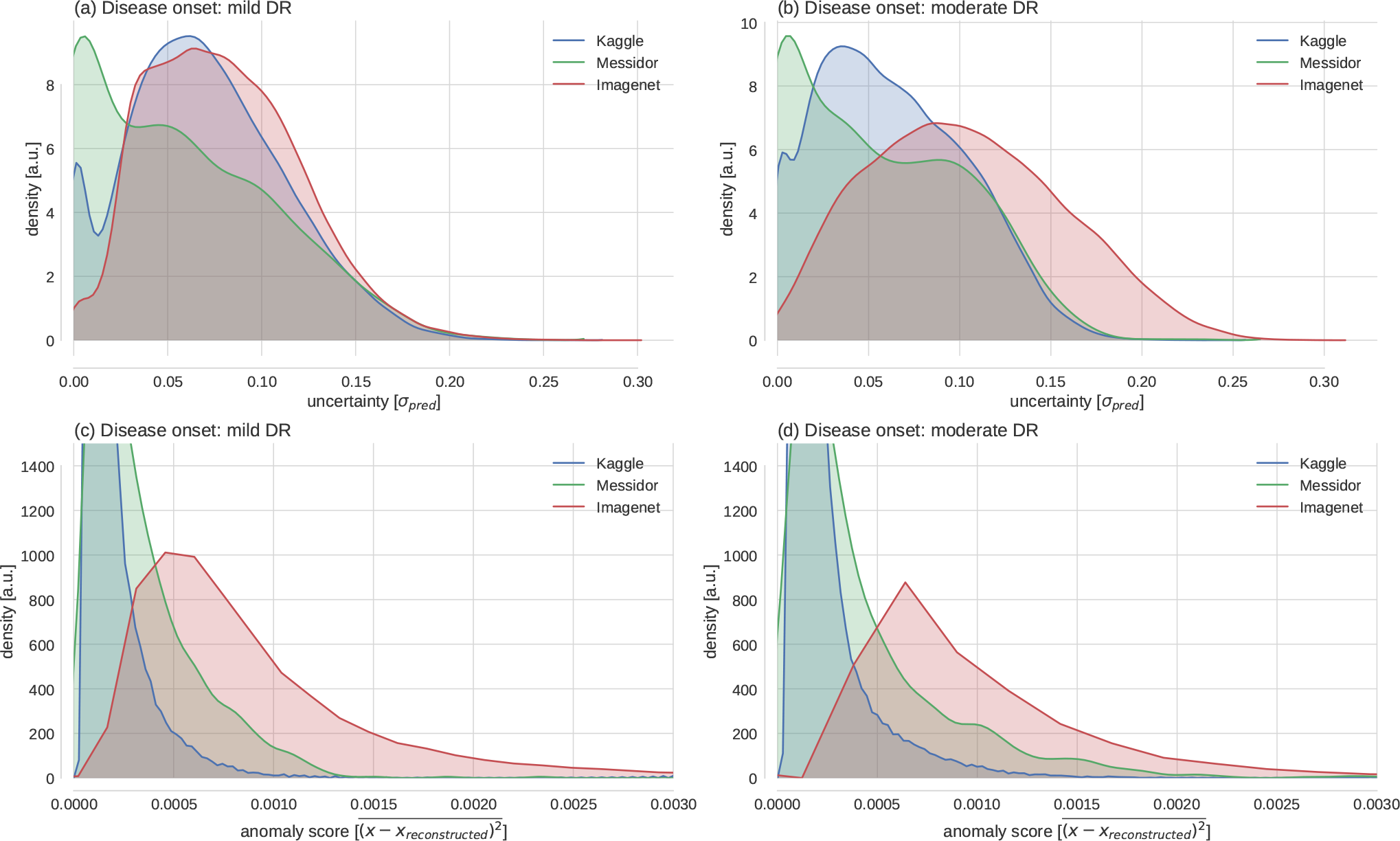
Uncertainty in face of (un)familiar data samples. (**a**) Empirical distributions of model uncertainty (σ_*pred*_) for familiar data with known semantic content (Kaggle) and unfamiliar data with known semantics (Messidor) vs. unknown semantics (Imagenet). (**b**) Same as in (a) but for the task of detecting moderate (2) instead of mild DR (1). (**c**) & (**d**) Reconstruction errors of a deep autoencoder trained on the penultimate layer features of the Kaggle training set. All distributions shown in a-d for Kaggle refer to the test set.

If the task is to detect samples far from the training distribution, an alternative to using the model uncertainty from the task dependent, discriminative setting (fig. 8 a,b) is to model the training data distribution in order to detect outliers. For comparison we therefore performed anomaly detection by autoencoding the 512-dimensional feature vectors of the penultimate layer of a disease detection network. The deep autoencoder (DAE) comprised two fully-connected (FC) encoding layers with 128 and 32 units, followed by two FC decoding layers with 128 and 512 units respectively. We quantified the distance of data samples from the (Kaggle) training data distribution by measuring the reconstruction error between a penultimate layer feature vector and its autoencoder based reconstruction (fig. 8c/d: disease onset 1/2). Here, Imagenet samples clearly show a higher anomaly score such that their presence could be detected based on the features learned by our network. The distribution of Messidor and Kaggle images seems similar, with Messidor images having slightly higher anomaly scores, being indicative of the fact that Kaggle and Messidor datasets have slightly different statistics.

## Discussion

Here we showed that it is feasible to associate deep learning based predictions about the presence of DR in fundus photographs with uncertainty measures that are informative and interpretable. Using the recently identified connection between dropout networks and approximate Bayesian inference^31,32^, we computed meaningful uncertainty measures without needing additional labels for an explicit *uncertain* category. Computing this uncertainty measure was efficient, as computing the approximate predictive posterior for one image took about ≈ 200*ms*. Seemingly appealing alternative uncertainty measures derived from either the conventional softmax output or GP classification applied to the penultimate layer features of our Bayesian networks were not found to be competitive for the purpose of deep learning based disease detection.

While not being crucially necessary for the purpose of evaluating model uncertainty, the performance achieved by our networks met the requirements for UK national standards for automatically diagnosing DR under several settings (Table 1, fig. 4). For all settings we could improve performance in terms of ROC AUC (fig. 4, S1) by referring uncertain ( fig. 5, 6, 7) cases for further inspection, outperforming alternative uncertainty measures throughout (fig., 4). Acquiring further opinions naturally integrates into the traditional clinical work flow as well as into a human-machine loop in which especially attended, difficult cases could be fed back into the model for its continuous improvement^59^.

We observed slightly worse performance on Kaggle data as compared to Messidor. We want to point out, that the quality of the former dataset was questioned previously - albeit informally, both by competition participants^47^ as well as by clinicians^54^. The extent to which the set of images considered uncertain by our approach overlaps with the images considered ungradable or wrongly labelled by humans is, however, unclear. Because images considered ungradable by clinical specialists may coincide with difficult diagnostic cases, these should be identifiable via high uncertainties from our approach. Easy decisions for images with wrong labels in turn should cause wrong predictions with low uncertainty. Both situations could hence be identified by our approach and be used to selectively reduce label noise and improve model performance.

The scope of which scenarios the assessed uncertainty is able to deal with can be understood by our results regarding the causes of uncertainty. We showed that model uncertainty was sensitive to clinically relevant cases, that is patients with an undefined disease state as determined by physicians (fig. 7). Aiming for a qualitative understanding, we showed that it is in particular difficult decisions that are considered uncertain by the networks, both for the 2D toy examples (fig. 5) as well as for the high-dimensional image case (fig. 6 & 7). The main difference of a Bayesian approach with respect to the plain softmax output is the fact that multiple separating hyperplanes instead of just a single one are taken into account. This renders the model uncertainty to extend beyond regions around the prediction uncertainty of the conventional network output (fig. 5a vs. c). Because we observed monotonic performance improvements under decision referral (fig. 4) and a composition of disease levels resembling the prior for intermediate uncertainties (fig. 6), a much higher fraction of the input space is associated with uncertainty (see as well fig. 8a,b) than the 2D scenario might suggest (fig. 5c). Recent research on uncertainty measures^18,34^is actually relying on this in order to detect unknown classes or adversarial attacks. The task dependency of discriminative model uncertainty together with high uncertainty due to diffuse class boundaries may however obscure the detectability of unfamiliar data samples (fig. 8a,b). Out-of-sample image detection was however feasible to some extent by modelling the data distribution in the penultimate layer’s feature space (fig. 8c,d).

We conclude that this work successfully demonstrated the benefits, applicability and limitations of uncertainty in deep learning^60^ for disease detection. This paradigm can readily be applied to recently published high performance disease detection networks^14,15^as well as other medical tasks and datasets as initial work on image registration^61^ and genome data^62^ has already shown. We also believe that segmentation^30^ and regression^63^ problems which are omnipresent in biomedical imaging and diagnostics could largely benefit from taking uncertainty into account.

## Methods

### General DNN methodology

#### Image preprocessing

All images were cropped to a squared centre region and resized to 512×512 pixels. In order to compensate for the decreased network depth in case of the Bayesian CNNs we additionally subtracted the local average colour for contrast enhancement purposes as described^67^ and used^13^ previously. Images fed to the JFnet were normalized the same way as had been used for training by the author^49^, whereas those fed to the BCNNs were standard normalized for each colour channel separately.

#### Network training

We trained one Bayesian CNN for each disease detection task using 80% of the Kaggle DR training data. We minimized the cross-entropy plus regularization terms (Eq. 5) using stochastic gradient descent with a batch size of 32 and Nesterov updates (momentum=0.9). All parameters were initialized with the weights from the JFnet. Final weights were chosen based on the best ROC AUC achieved on a separate validation set (20% of Kaggle DR training data) within 30 training epochs. The learning rate schedule was piecewise constant (epoch 1-10: 0.005, epoch 11-20: 0.001, epoch 21-25: 0.0005, epoch 26-30: 0.0001). L2-regularization (*λ* = 0.001) was applied to all parameters, L1-regularization (*λ* = 0.001) to only the last layer in the network. Data augmentation was applied to 50% of the data in an epoch. Affine transformations were composed by drawing uniformly from ranges for zooming (±10%), translating (independent shifts in x-and y-directions by ±25 pixels), and rotating (±*π*). Transformed images were in addition flipped along the vertical and/or the horizontal axis if indicated by respective draws from a Binomial distribution (*µ* = 0.5). Effects of class imbalance onto the stochastic gradient were compensated by attributing more weight to the minority class, given by the relative class frequencies in each mini-batch^68^ *p(k)*_*mini-batch*_. To achieve this, we reweighed the cross-entropy part of the cost function (compare Eq. 5) for a mini-batch of size *n* to:

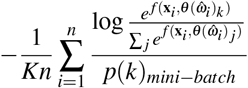

We fixed the amount of dropout for the convolutional layers to *p*_*drop*_ = 0.2, because this was a good compromise between getting a reasonable performance and uncertainty measures. We observed convergence problems for larger *p*_*drop*_ when initializing the Bayesian CNNs with the pretrained weights from the network without dropout between conv. layers. Gradually increasing dropout during training could potentially ease convergence. Alternatively, the dropout rates could be learnt via *variational dropout*^28^.

### Approximate Bayesian model uncertainty for deep learning

Recently, it was shown^31^ that a multi-layer-perceptron (i.e. a stack of densely connected layers) with dropout after every weight layer is mathematically equivalent to approximate variational inference^52^ in the deep GP model^69,70^. This result holds for any number of layers and arbitrary non-linearities. Next, this idea was extended to incorporate convolutional layers^32^, potentially loosing the GP interpretation, but preserving the possibility to obtain an approximation to the predictive posterior in a Bayesian sense. Here, we summarize the core idea for deep classification networks and highlight in particular the difference between the Bayesian perspective and the classification confidence obtained from the softmax output.

#### Softmax vs. Bayesian uncertainty

DNNs (with or without convolutional layers) for classifying a set of *N* observations {**x**_1_,…, **x**_*i*_,…, **x**_*N*_} into a set of associated class memberships {**y**_1_,…,**y**_*i*_,…,**y**_*N*_} with y_*i*_ ϵ {1,…,*K*}, and *K* the number of classes, can be trained by minimizing the cross-entropy between the distribution of the true class labels and the softmax network output:

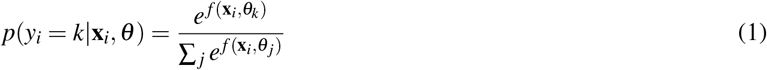

Equation (1) denotes the probability that the observation x_*i*_ belongs to class *k*, if propagated through the network function *f* with all parameters summarized by *θ*, i.e. weights **W**_*i*_ and biases **b**_*i*_ of all layers *i* ϵ {1,…,*L*}. For the example of disease detection from images, we have a single unit whose output denotes the probability for the presence of the disease in a given image.

Cross-entropy minimization results in a single best parameter vector *θ*, constituting the maximum-likelihood solution. *L*2-regularization, typically used to prevent overfitting, is equivalent to putting a Gaussian prior on the network parameters, resulting in a maximum-a-posteriori (MAP) solution.

A fully probabilistic treatment in a Bayesian sense, however, would consider a distribution over network parameters instead of a point estimate. By integrating over the posterior *p*(*θ* |**X**, **y**, **x**\*) given the entire training data {**X**, **y**} and a new test sample **x**\* one would like to obtain the predictive posterior distribution over class membership probabilities:

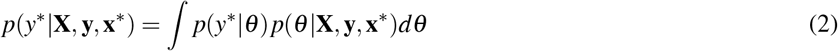

Whereas equation (1) determined a single value specifying the probability that an image belongs to the *diseased* class, the predictive posterior (Eq. 2) defines a distribution of such predictions, that is the probability values that a single image is *diseased*. Intuitively, the width of the predictive posterior should reflect the reliability of the predictions. For large training data sets, the parameter point estimates (from maximum-likelihood or MAP) may correspond to the mean or mode of the predictive posterior, resulting in a potentially strong relationship between the width of the predictive posterior and the softmax output, however this is not guaranteed. Indeed we’ve found that only for the original JFnet the softmax output may be used as a proxy for (prediction instead of model) uncertainty (values close to 0.5 were considered uncertain), whereas the Bayesian treatment worked for all investigated scenarios.

#### Bayesian convolutional neural networks with Bernoulli approximate variational inference

In practice, equation (2) is intractable and a common way to find approximating solutions is via *variational inference.* We assume the true posterior to be expressible in terms of a finite set of random variables *ω*. The posterior is then approximated by the *variational distribution q(ω)* as follows:

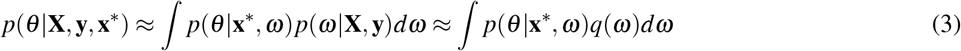

Maximizing the *log evidence lower bound* with respect to the approximating distribution *q(ω)*:

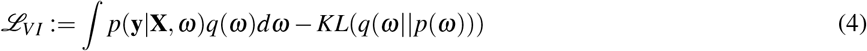

has two effects. The first term maximizes the likelihood of the training data {**X**, **y**}, whereas the second term takes care of approximating the true distribution *p(ω)* by *q*(*ω*). The key insight from *Gal & Ghahramani* was then to link equation (4) with dropout training. Here, we will summarize the derivations^35^ in words. The integral in eq. (4) is still intractable and therefore approximated with Monte Carlo sampling. This results in the conventional softmax loss for dropout networks, for which units are dropped by drawing from a Bernoulli prior with probability *p*_*drop*_ for setting a unit to zero. The KL term in (4) was shown^35^ to correspond to a L2-regularization term in dropout networks. Summing up, approximate variational inference with a Bernoulli approximating distribution can be performed with the following loss:

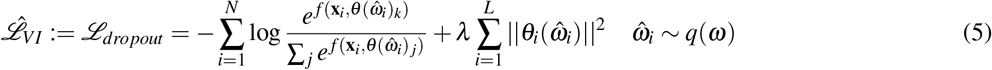

We use 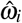 as a shorthand notation for stating that in order to decide whether a unit is dropped, we independently sample from a Bernoulli distribution (with probability *p*_*drop*_) for each unit in all layers for each training sample. Note that Monte Carlo sampling from *q(ω)* is equivalent to performing dropout during training, hence we get the Bayesian network perspective as well for already trained models.

#### Obtaining model uncertainty at test time

Obtaining model uncertainty for a given image is as simple as keeping the dropout mechanism switched on at test time and performing multiple predictions. The width of the distribution of predictions is then a reasonable proxy for the model uncertainty. More formally expressed, we replace the posterior with the approximating distribution Eq. 3 and plug it into the expression for the predictive posterior (2):

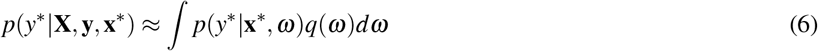

We then approximate the integral by Monte Carlo sampling and compute the predictive mean (to be used for a final prediction on a test image):

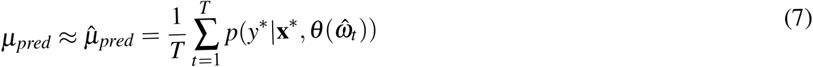

as well as the predictive standard deviation as a proxy for the uncertainty associated with this prediction:

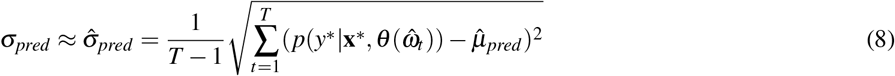

An intuitive illustration is given in the main text (fig. 1). In short: the wider the distribution, the less the different subnetworks agree. Regarding the choice of *T*, we fixed it to *T* = 100 because it was shown by^32^ to suffice for reducing the test error by improving the quality of the predictive mean estimation. Different values for *T* affect how well the true population moments µ_*pred*_ and *σ*_*pred*_ can be estimated: 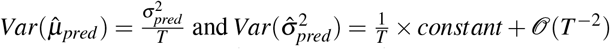^71^. It should however be noted, that the computation of MC samples is extremely fast: The test predictions could be performed in parallel, but even a serial implementation takes less than 200*ms* per image. Hence there is no practical reason to compute fewer testing samples than the *T* = 100 used to obtain the results presented here.

#### Analysis of results

All density plots are based on Gaussian kernel density estimates, for which the bandwidth was chosen based on Scott’s method^72^. All line plots are based on the entire data and the 95% confidence intervals were obtained from 10^4^ bootstrap samples.

### Data and code availability

We used the deep learning framework Theano^64^(0.9.0dev1.dev-RELEASE) together with the libraries Lasagne^65^(0.2.dev1) and Keras^66^(1.0.7). Network trainings and predictions were performed using a NVIDIA GeForce GTX 970 and a GeForce GTX 1080 with cuda versions 7.5/8 and cuDNN 4/5. For the GP analysis, we used the Gaussian Processes for Machine Learning (GPML) Toolbox^73^ (v3.6). All code and models for fast DR detection under uncertainty will be publicly available upon publication at https://bitbucket.org/cleibig/disease-detection.

## Acknowledgements

This work was funded by the German excellence initiative through the Institutional Strategy of the University of Tubingen and the Center for Integrative Neuroscience (EXC 307), the Bernstein Award for Computational Neuroscience by German Ministry for Education and Research (BMBF; FKZ: 01GQ1601) to PB. Additional support came from the early career program of the Medical Faculty of the University of Tubingen.

## Author contributions statement

C.L., P.B. and S.W. designed the concept of the study. C.L. conducted the study. M.S.A. conducted the GP analysis and wrote respective sections. C.L., V.A. and P.B. wrote the manuscript. All authors reviewed the manuscript.

